# Community diversity and interaction of bacterial, cyanobacteria and protist among aquatic and soil micro-food chains

**DOI:** 10.64898/2026.07.31.741965

**Authors:** Tiantian Sun, Xinke Yu, Lina Wei, Shanmei Zou

## Abstract

Microorganisms, found in every environment, play an important role in recycling matter and providing energy to the ecosystem. Here, we used 18S and 16S metabarcoding to study bacteria, cyanobacteria and protist community diversity and functional ecology among different environments of typical lake, river, marine and soil from the Yangtze Delta of China. The results showed that the similarity of cyanobacteria and protozoa communities in soil and aquatic environments was both higher than that of bacteria and microalgae communities. While the α diversity of cyanobacteria and bacteria in lake was higher than that in river, marine and soil, the α diversity of microalgae and protozoa in river was higher than that in lake, marine and soil. The β diversity of cyanobacteria and bacteria marked differences in lake from other environments. But The distribution of dominant families of cyanobacteria, bacteria, microalgae and protozoa is similar to α diversity. While significant positive correlations were found among dominant species of cyanobacteria, protozoa, bacteria and microalgae in lake the dominant species among bacteria, microalgae and protozoa in river, marine and soil all showed more negative correlations. Our study provides the basis for understanding the functional ecology of microbes in the micro-food webs of different environments.

## Introduction

Yangtze River Delta has paid a huge price for the rapid urbanization of the region. It has become a new ecological fragile area in China (Wang et al. 2015).On the one hand, water pollution, acid rain, soil pollution, solid waste accumulation and other problems are becoming increasingly prominent, and cross-border pollution problems continue to cause inter-regional conflicts (Gu et al., 2011). The Beijing-Hangzhou Grand Canal Yangtze River Delta, Taihu Lake, the lower reaches of the Yangtze River and Qiantang River are all polluted to varying degrees, among which Taihu Lake is the most polluted (Guan et al. 2011). In recent decades, lake-wide cyanobacteria blooms have occurred every year in Taihu Lake, destroying the natural function of the lake and threatening the safety of drinking water resources (Fu et al. 2015; Pan et al. 2006; Ye et al. 2011). Nanjing is one of the most economically developed cities in China, located in the middle and lower reaches of the Yangtze River, with complex industrial structure (J Wang et al. 2016), rapid urbanization and high utilization rate of soil agriculture (H Wang et al. 2018). However, with the rapid development of industry and urban modernization, ecological environment problems are becoming increasingly prominent (C Wang et al. 2015).

Microorganisms are present in every habitat on earth, like soil, water, animal, and plants as well as humans. Both prokaryotic and eukaryotic microbes in earth play key functional roles in all ecosystems, particularly by catalyzing carbon and nutrient cycling (Stahl et al. 2012). The prokaryotic microbes mainly include bacteria and cyanobacteria while the eukaryotic microbes include microalgae and protozoa. Although a substantial proportion of microbes has been described (Archibald et al. 2017) the right order of magnitude is unknown (Whitfield 2005). However, estimating both prokaryotic and eukaryotic microbes is always challenging because of their tiny body and similar external shapes. With the high-throughput sequencing, the environmental DNA-based methods have revealed a high and largely unknown taxonomic diversity of both prokaryotic (Delgado-Baquerizo et al. 2018) and eukaryotic microbes (de Vargas et al. 1999; Tedersoo et al. 2014).

Microalgae and cyanobacteria are primary producers in the food chain of different environments because they fix carbon into organic compounds through photosynthesis and thus provide food sources for all other organisms, including protozoa, within a food web (OR Anderson 2001; Singer et al. 2021; Tortora et al. 2004). Heterotrophic protist, particularly the protozoa, are consumers within food webs since they prey on algae and other microbes (OR Anderson 2001). Protozoa are also often used as bioindicators of environmental pollution (OR Anderson 2001). Bacteria and fungi can primarily biodegrade hydrocarbons in the environment (Das et al. 2011; Leahy et al. 1990). In aquatic ecosystem, bacteria utilize organic compounds released by primary producers, or by death and decay of other biota, to form a major particulate food source for small and intermediate-sized heterotrophic protozoa, resulting in replenishment of available nutrient resources (Fenchel 2008). In soil environment, food webs are largely bacterial-based and productivity depends on the abundance and suitability of bacteria as prey (DM Anderson et al. 2021) while the soil microalgae and cyanobacteria release nutritive products, organic products and active components that can serve as food for bacteria and invertebrates (Abinandan et al. 2019).

The ecological environment of the microbial biodiversity in freshwater, marine and soil differs much due to their various environmental habitats, including salinity, average temperature, depth, and nutrient content (Abinandan et al. 2019; Aryal et al. 2015; Singer et al. 2021). Assessing how they vary among taxonomic, biomes and ecosystems initiates the understanding of their ecosystem functioning and associated services (Endo et al. 2020), which is still relevant question in biology (Mora et al. 2011). Singer et al (2021) presented a comparison of protist diversity based on standardized high throughput 18S rRNA gene sequencing of soil, freshwater and marine environmental DNA. To date, however, a comprehensive comparative analysis of diversity, interaction and functional ecology of bacteria, microalgae and protozoan among soil and aquatic system is missing. Here we employ metabarcoding from 16S, 16S-v3v4 and 18S-v4 genes to compare the diversity and interaction of bacteria, cyanobacteria, microalgae and protozoa community from typical soil, river, lake and marine environments of Yangtze Delta of China, and evaluate the functional ecology within the micro-food webs. It provides theoretical reference for biological governance of different environments in the Yangtze River Delta region.

## Material and Methods

### Sample collection

The samples were collected from park soil in Nanjing city, the Yangtze River, Tailake and Shengsi marine region of Yangtze Delta of China (Fig. 1). Water samples were firstly pre-filtered through a 200-μm pore-size sieve to remove debris, mesoplankton, and macroplankton. Then each water sample (∼500 ml) with bacteria, cyanobacteria and microeukaryotes was subsequently filtered through a 0.2-μm pore-size polycarbonate membrane (Millipore, Billerica, MA, USA). Soil samples measuring 10 m × 10 m were selected from each site. In order to avoid cross-contamination during sampling process, the sampling tools should be disinfected with alcohol for each sampling point. All the processed samples of various environments were stored in −80℃until DNA extraction. In total, 20 samples were collected for each of soil, river, lake and marine environments.

**Figure. 1.**
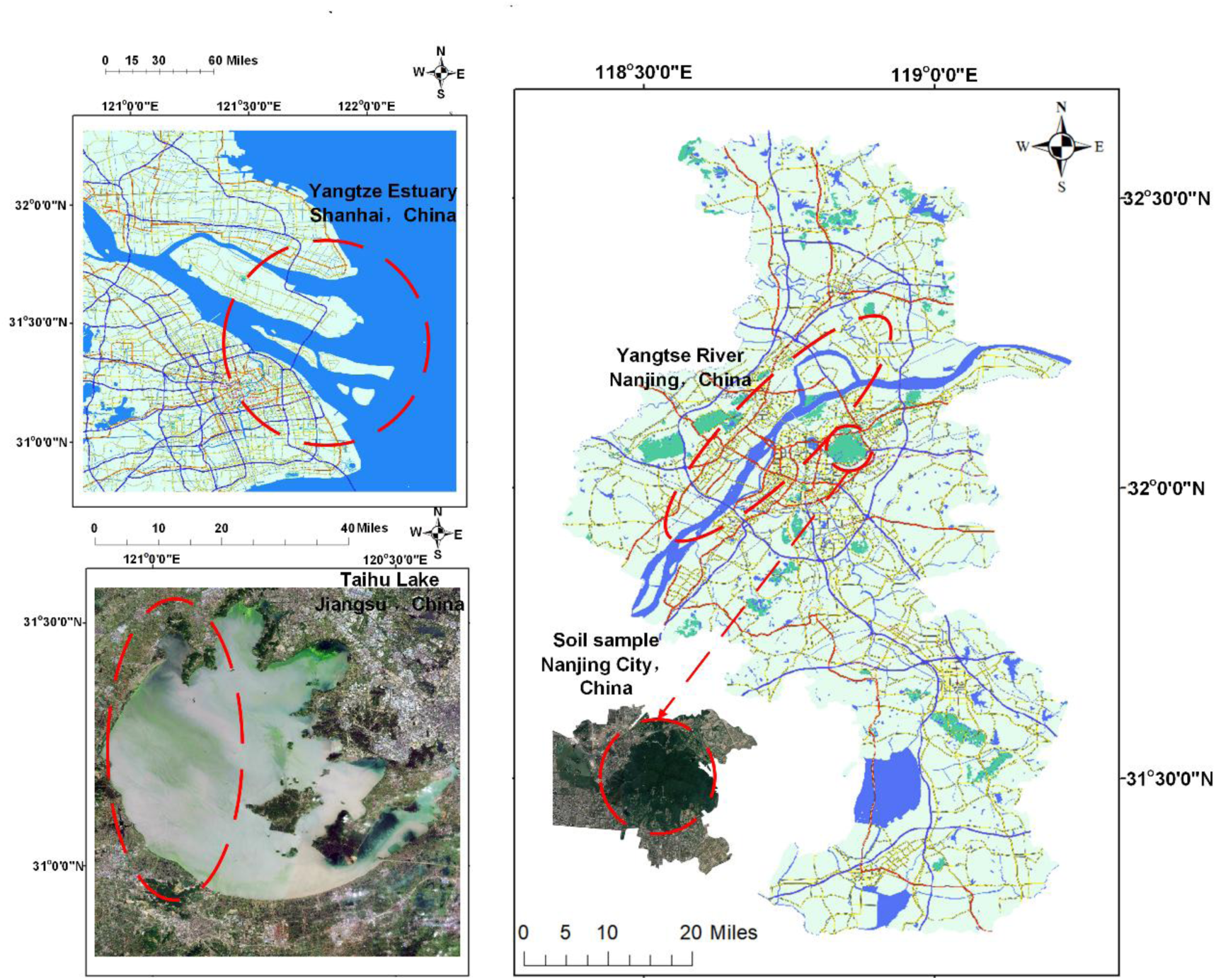
Marine, lake, river and soil sample collection areas.

### DNA extraction, PCR amplification and sequencing

DNA was extracted by OMEGA E.Z.N.ATM Mag-Bind Soil DNA Kit for each sample. The PCR amplification of all samples were triple. The 16S, 16S-v3v4 and 18S-v4 genes were amplified for cyanobacteria, bacteria and microeukaryotes respectively (Table 1). Primers were synthesized with the locus-specific sequence on the 3’ end and a 5’ tail containing sequences matching the TruSeq sequencing primer binding site. Triple PCR products for each sample were pooled, which was then cleaned by Solid Phase Reversible Immobilization (SPRI) Beads. Finally, all the amplicons were pooled in equal amounts for sequencing on an Illumina MiSeq 600 cycles.

**Table 1.**
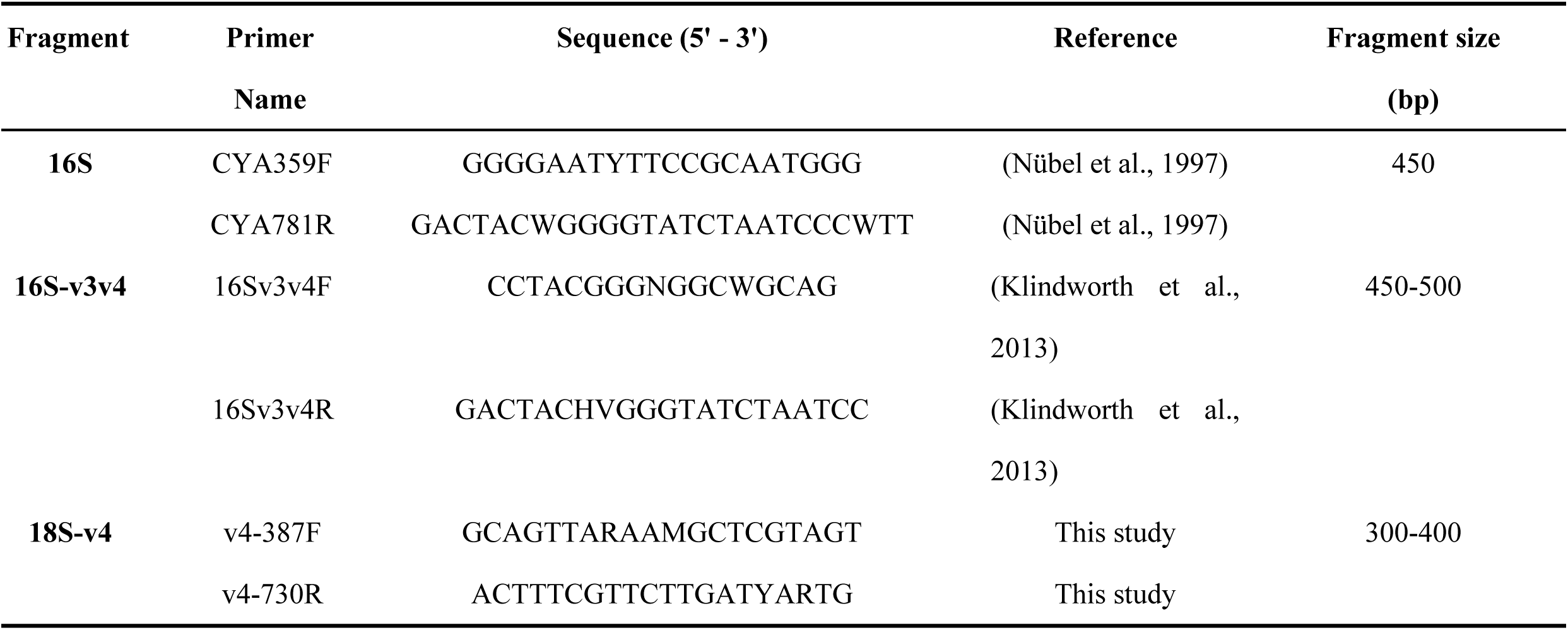
Multiplex PCR systems and primers used in this study.

### Bioinformatics analysis

The reference database for each genetic marker were downloaded from EMBL standard nucleotide database by ecoPCR (Ficetola et al. 2010). Raw sequence fastq files were firstly demultiplexed to separate the samples based on the indexes. Then all sequence analysis was performed using QIIME2 (Bolyen et al. 2019). The PCR primers were removed by cut adapt package. Dada2 was applied for denoising, dereplicate-sequence, filtering chimera and merging the paired reads. Taxonomic classification was performed by feature-classifier module with Naive Bayes. We filtered the ASVs with assignment score under 0.75 for all genes.

### Statistics

The α diversity was calculated using QIIME2 to evaluate the richness and diversity. We assessed the similarity patterns among microbial communities of different environments (β-diversity) by non-metric multidimensional scaling (NMDS). NMDS was calculated on Bray-Curtis dissimilarities retrieved from the sequence relative abundance. All statistical analysis were performed in the R program (Pinheiro et al. 2016) using the packages vegan (Oksanen et al. 2019). The Perl package Circos (version 0.64) (Krzywinski et al. 2009) was used for numerical visualization of positive and negative correlations among different taxonomic groups from different environemts.

## Results

### Taxonomic composition and abundance

The mean relative abundance of each taxonomic group in each environment was presented (Fig. 2). In lake, the cyanobacteria and bacteria accounted for considerable proportion. Cyanobacteria also dominated in soil. In river, the protozoa and microalgae were dominant groups. The cyanobacteria and protozoa accounted for higher proportion in soil.

**Figure. 2.**
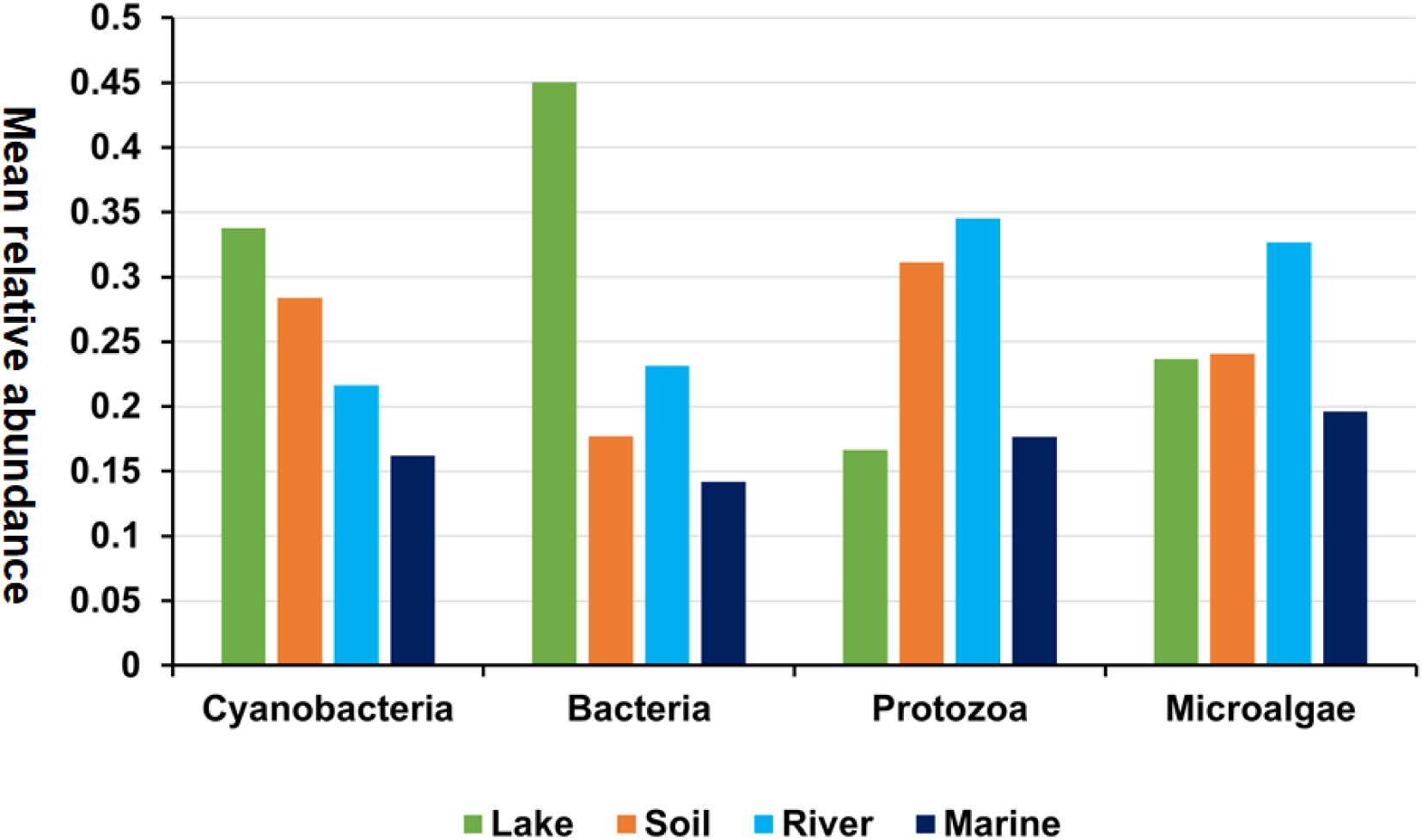
Average relative abundance ratio of different microbial communities in different environments.

For cyanobacteria at order level, both *Synechococcales* and *Oscillatoriales* were found in four environments (Fig. 3A). *Chroococcales* showed the highest abundance in lake while *Oscillatoriales* showed the highest abundance in soil and marine environments. *Nostocales* had the highest abundance in river. For bacteria, at class level (Fig. 3B), *Alphaproteobacteria* and *Actinobacteria* existed in all four environments. While *Alphaproteobacteria* had the highest abundance in lake, soil and marine *Actinobacteria* had the highest abundance in river. For Protozoa, at the order level, *Bicosoecida*, *Cercomonadida*, *Sessilida* and *Haptorida* were found in all four environments, with *Bicosoecida* having the highest abundance in rive and marine environments (Fig. 4A). Also, while *Cryomonadida* had the highest abundance in soil *Sporadotrichida* had the highest abundance in lake (Fig. 4A). For microalgae, at the order level, *Chromulinales* existed in all four environments (Fig. 4B). *Thalassiosirales* had the highest abundance in lake and river, and *Naviculales* had the highest abundance in soil. *Gymnodiniales* had the highest abundance in river (Fig. 4B).

**Figure. 3.**
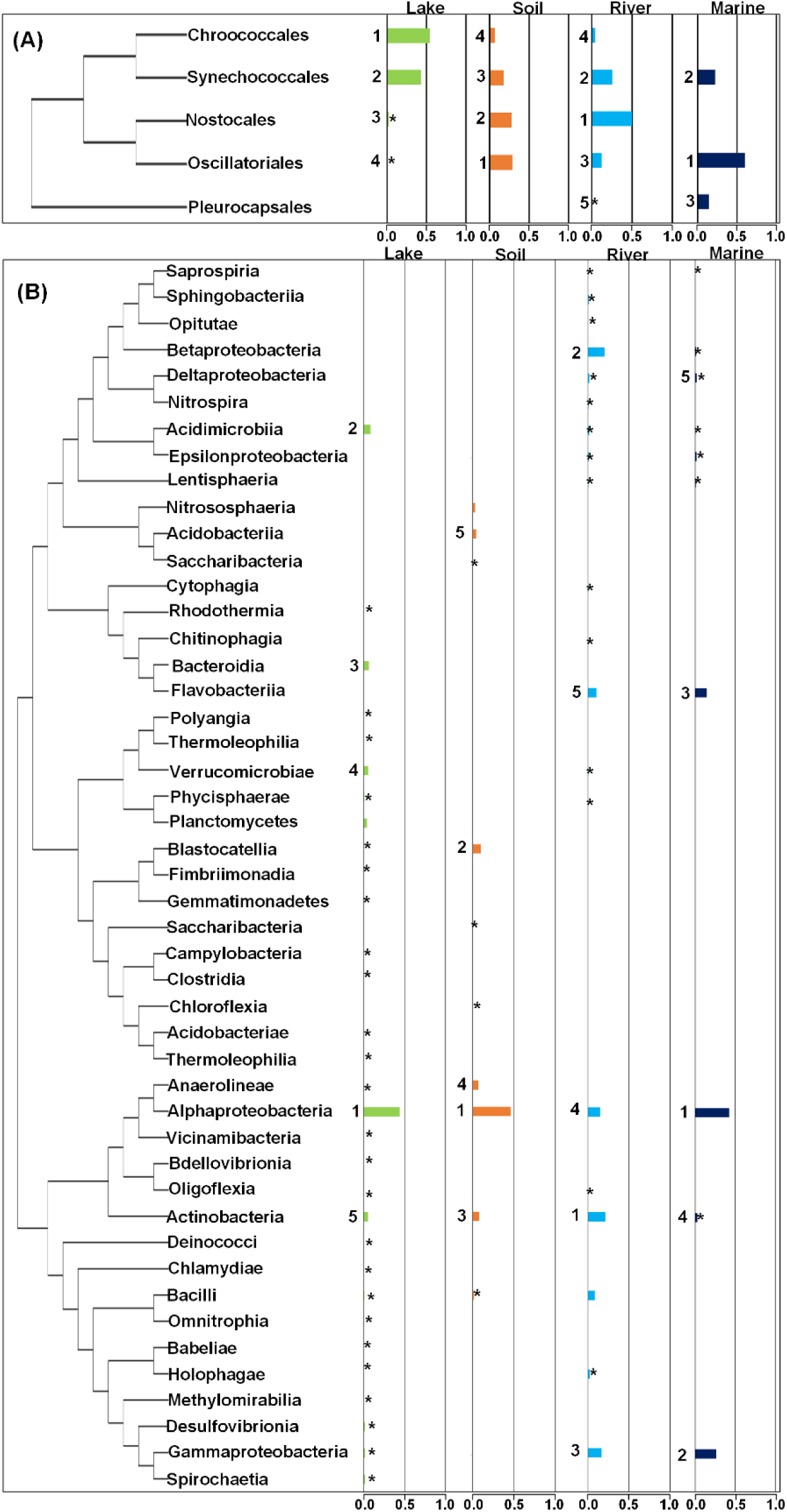
**A.** Schematic phylogenetic tree and their mean relative abundance of major groups of cyanobacteria (order) in four ecosystems. **B.** Schematic phylogenetic tree and their mean relative abundance of major groups of bacteria (class) in four ecosystems. Lakes: green, Marine: dark blue, River: cyan, Soil: orange. Numbers 1-5 represent the top5 of the distribution in their respective ecosystems. * Indicates an average relative abundance of less than 3%.

**Figure. 4.**
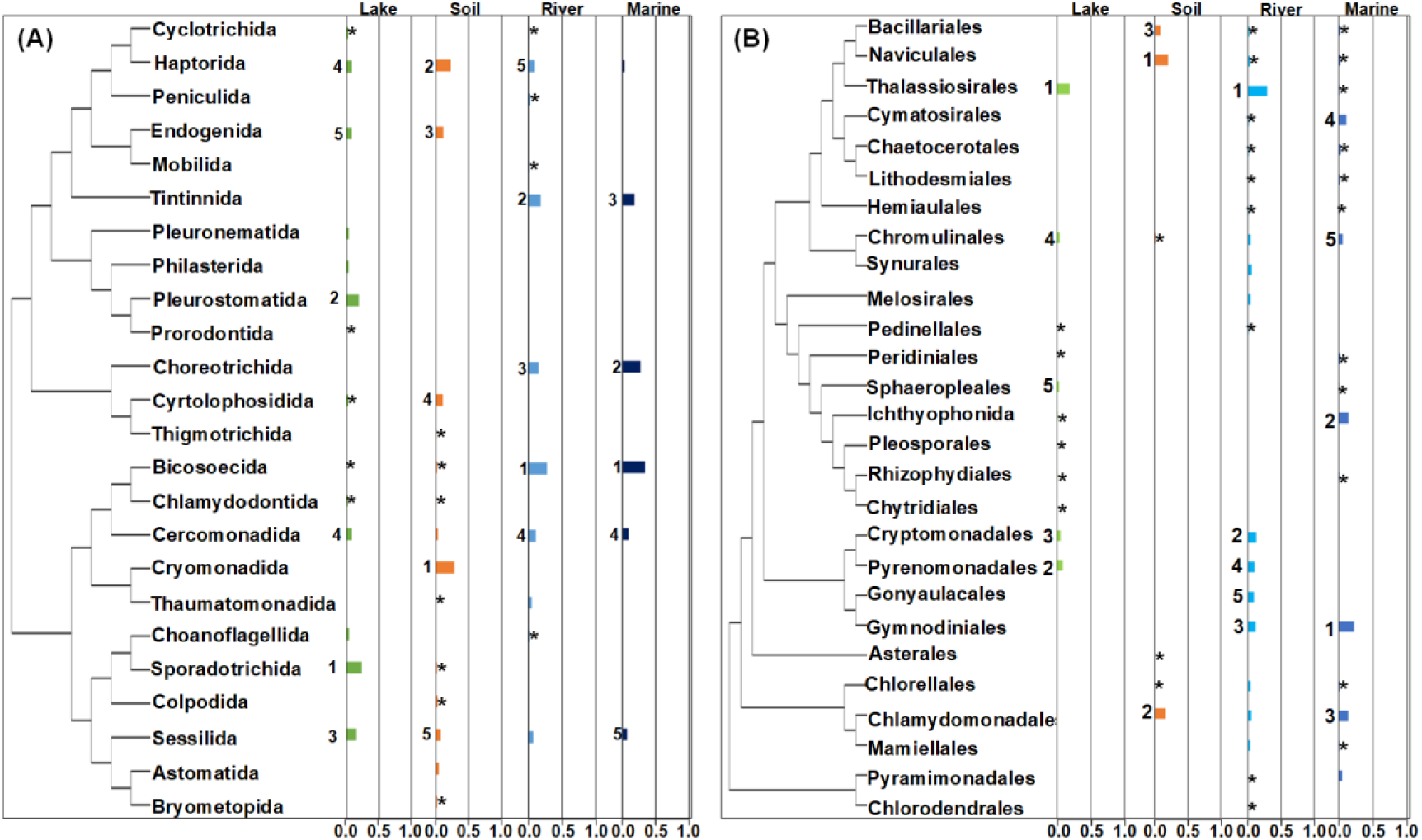
**A.** Schematic phylogenetic tree and their mean relative abundance of major groups of protozoa (order) in four ecosystems. **B.** Schematic phylogenetic tree and their mean relative abundance of major groups of eukaryotic microalgae (order) in four ecosystems. Lakes: green, Marine: dark blue, River: cyan, Soil: orange. Numbers 1-5 represent the top5 of the distribution in their respective ecosystems. * Indicates an average relative abundance of less than 3%.

### α and β diversity

For cyanobacteria, the richness index, Chao index and ACE index in lake environment were higher than that in other three environments (P<0.05), and the Shannon index, Simpson index and invsimpson index in river environment were higher than that in other three environments (P<0.05) (Fig. 5A). The richness index, Shannon index, Simpson index, invsimpson index and Chao index of bacterial communities in lake environment were higher than that in other three environments (P<0.05) (Fig. 5B). The richness index, Shannon index, Simpson index, invsimpson index, Chao index and ACE index of protozoa and microalgae were higher than that of the other three environments (P<0.05) (Fig. 5C and D). The Pielou index of cyanobacteria, bacteria, protozoa and eukaryotic microalgae in soil was higher than that in lake, river and marine environments (P<0.05) (Fig. 5).

**Figure. 5.**
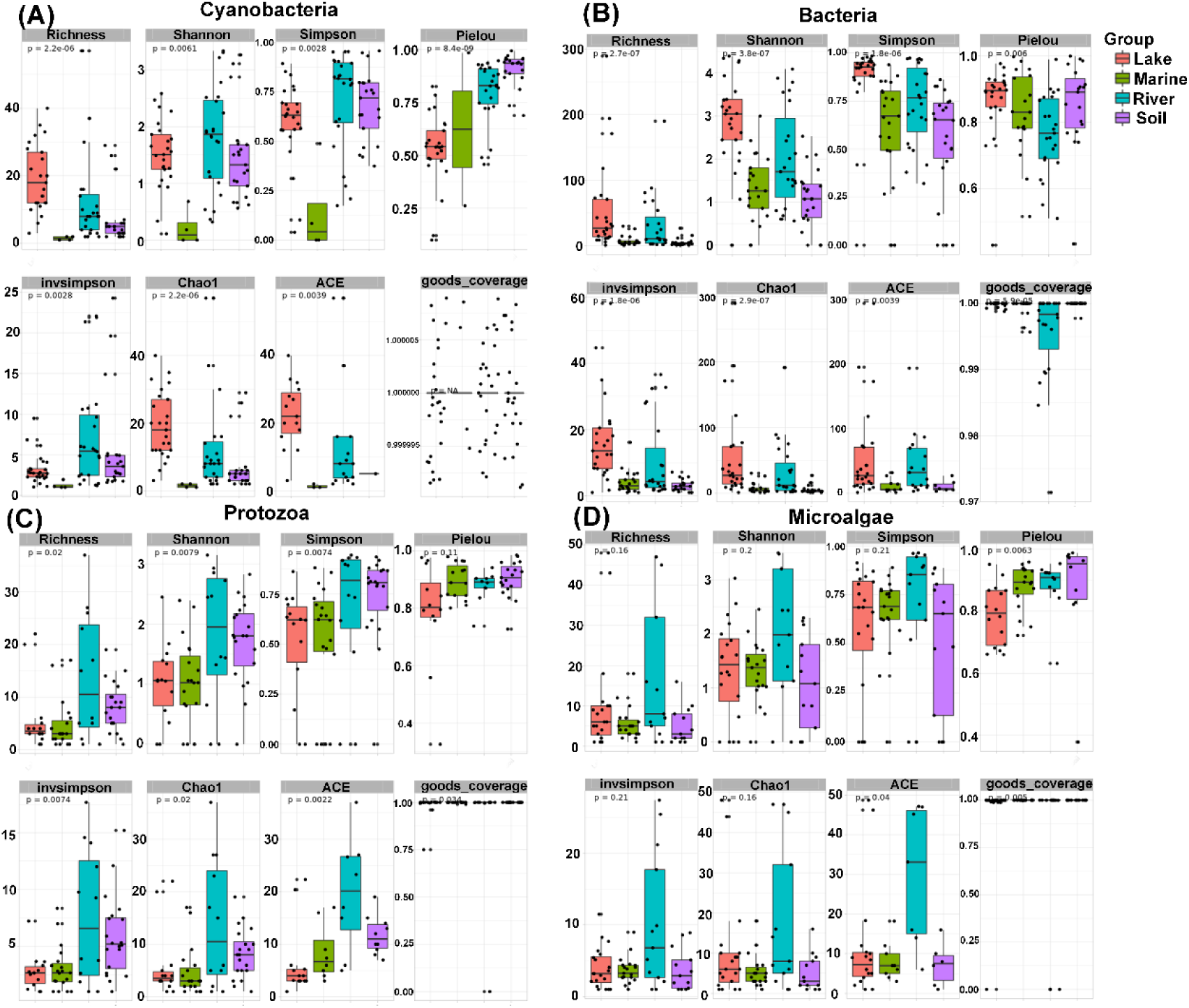
The α diversity index of cyanobacteria, bacteria, protozoa and eukaryotic microalgae communities was analyzed based on lake, marine, river and soil. **A.** cyanobacteria **B.** bacteria **C.** protozoa **D.** eukaryotic microalgae.

Based on Bray-Curtis similarity coefficient, NMDS analysis showed that the stress index of cyanobacteria, bacteria, protozoa and microalgae were all less than 0.2 (Fig. 6). Generally, the cyanobacteria and bacteria were distinct from other groups in the lake (Fig. 6A and B), and the cyanobacteria, bacteria, protozoa and microalgae in river, marine and soil were not separately clearly (Fig. 6).

**Figure. 6.**
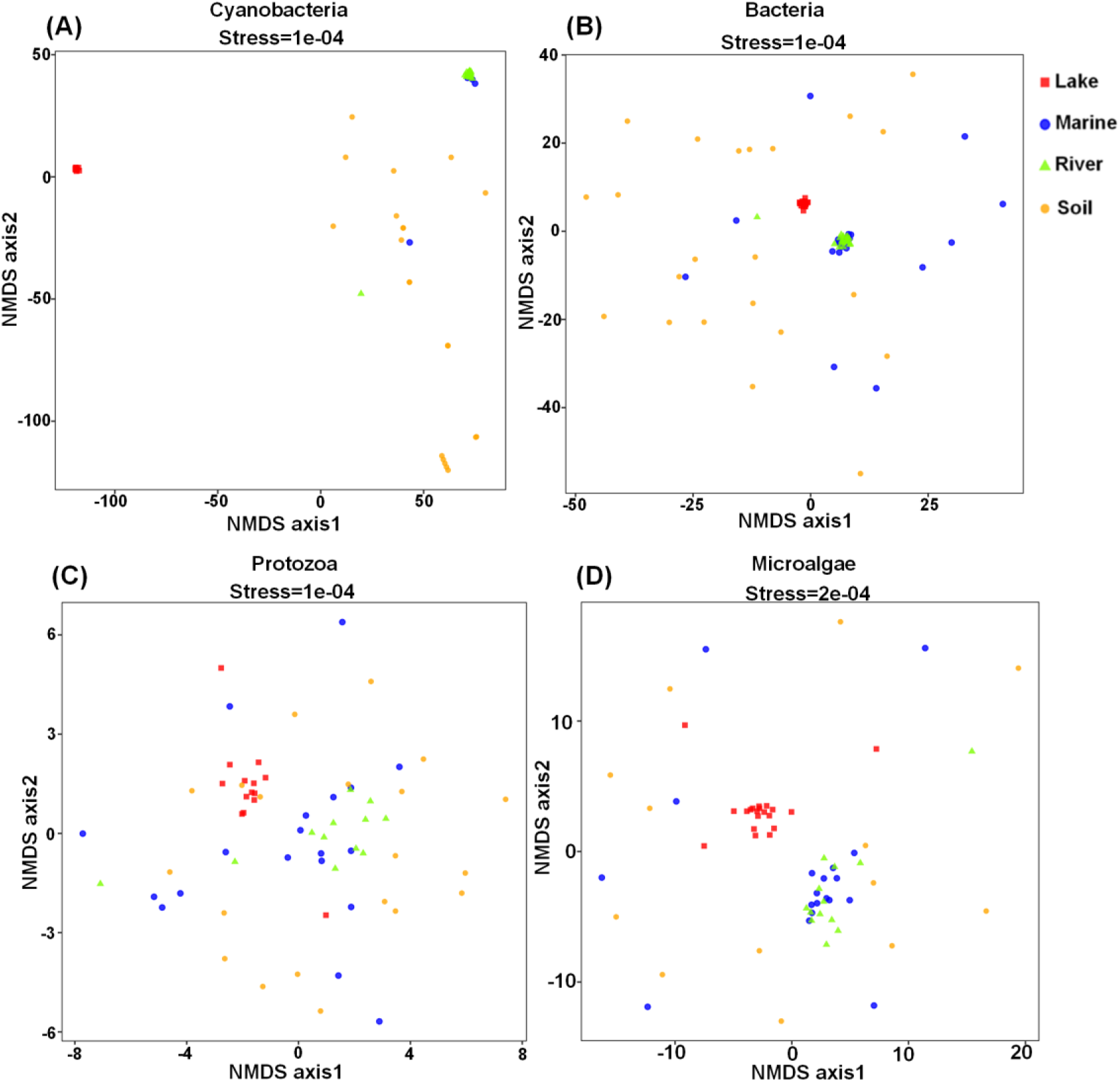
NMDS analysis of cyanobacteria, bacteria, protozoa and microalgae communities based on lake, marine, river and soil environments. **A.** cyanobacteria **B.** bacteria **C.** protozoa **D.** eukaryotic microalgae.

### Distribution of dominant families

The Circos chart showed the dominant 10 families of cyanobacteria, bacteria, protozoa and microalgae among four environments (Fig. 7). For cyanobacteria, Rivulariaceae and Synechococcaceae accounted for 62% and 39% of mean relative abundance respectively in lake (Fig. 7A). Rivulariaceae was also distributed in soil, and Aphanizomenonaceaez was also distributed in lake and river (Fig. 7A). For bacteria, the dominant families in the lake were Rhodobacteraceae and Ilumatobacteraceae. Rhodobacteraceae could also be found in lake, river and marine environments. While Flavobacteriaceae was dominant in lake, river and marine Sphingomonadaceae was dominant in soil (Fig. 7B). For microalgae protozoa, Stephanodiscaceae and Cryptomonadaceae accounted for 24% and 22% of mean relative abundance respectively in river (Fig. 7C). Brachysiraceae was also found in all of four environments (Fig. 7C). For protozoa, Codonellidae and Halteriidae were both found in lake with 29% of mean relative abundance. Strombidinopsidae and Tintinnidae accounted for 26% and 33% of mean relative abundance respectively in river. Strombidinopsidae also accounted for a high proportion of abundance in the marine, and Dileptidae was the dominant family in soil (Fig. 7D).

**Figure. 7.**
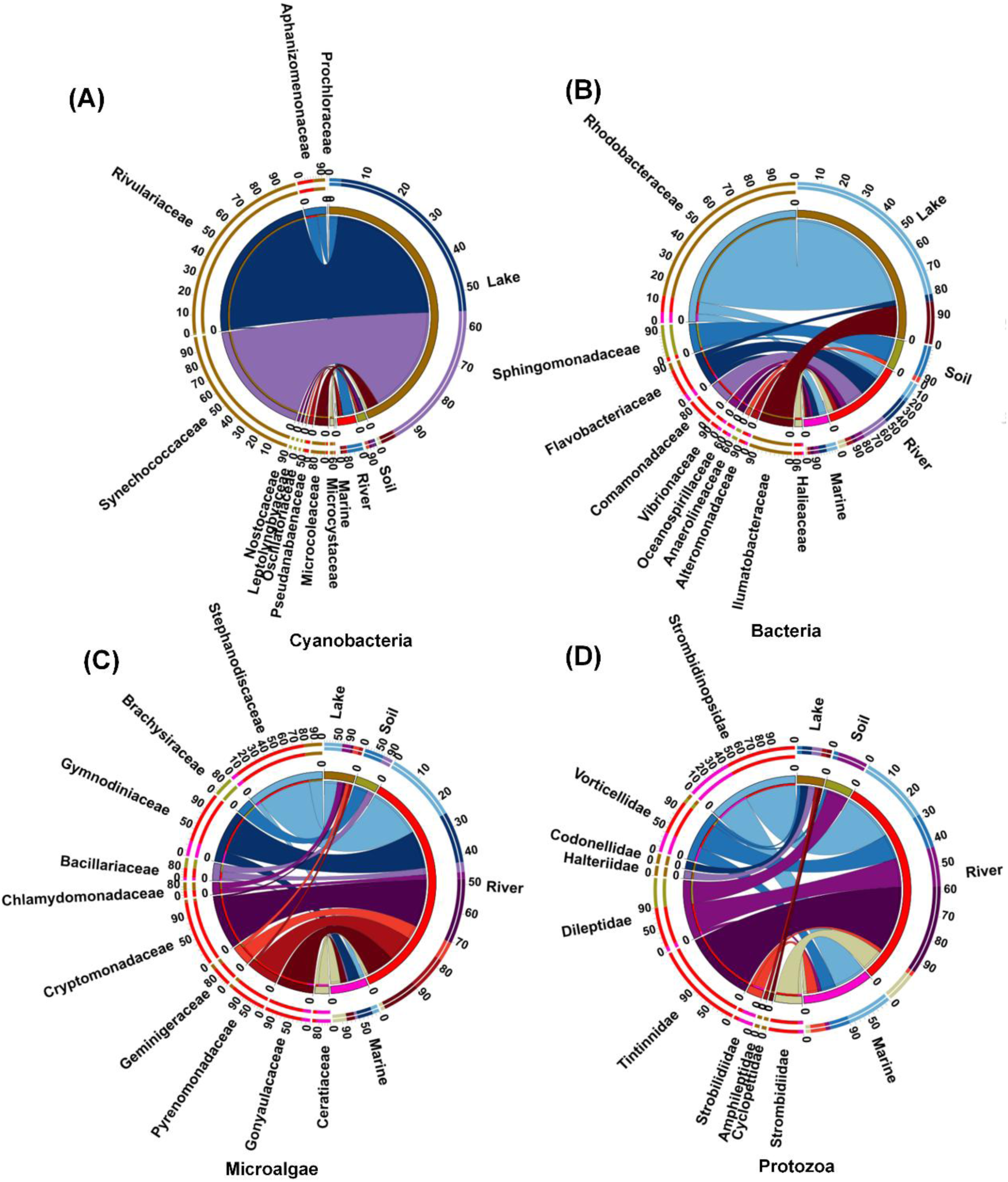
Distribution of cyanobacteria, bacteria, protozoa, and eukaryotic microalgae communities (families) in lake, river, marine, and soil. **A.** cyanobacteria **B.** bacteria **C.** protozoa **D.** eukaryotic microalgae. The small semicircle color on the left represents the composition of the Top10 dominant families, the outer circle number is the percentage representing the average relative abundance of the species, the small semicircle color on the right represents the distribution of the species in different environments, and the outer circle number is the percentage representing the proportion of the species in the environment.

### Correlation of dominant species

The dynamic Circos heat map showed the correlations among top 10 dominant species for each environment (Fig. 8, Fig. 9 and Fig. S1). Significant positive correlations were found among cyanobacteria, protozoa, bacteria and eukaryotic microalgae in lake (Fig. 8A, B and Fig. S1A). The *Microcystis* sp1 was the dominant cyanobacteria species with the highest mean relative abundance in lake, which was positively correlated with the dominant protozoa species *Halteria* sp1, *Pseudovorticella* sp1 and *Tintinnopsis* sp2, and was negatively correlated with the other dominant protozoa species (Fig. 8A). The *Microcystis* sp1 in lake was also positively correlated with the dominant bacteria species *Porphyromonas sp.*, *Terrimicrobium* sp2, *Rhodobacter* sp1 and *Rhodopirellula* sp1, and was negatively correlated with the other bacteria dominant species (Fig. 8B). The dominant microalgae species *Chlamydomonas* sp., *Stephanodiscus* sp1 and *Cyclotella choctawhatcheeana* were positively correlated with *Microcystis* sp1 (Fig. S1A). The other nine dominant cyanobacteria species were also mainly positively correlated with protozoa, bacteria and eukaryotic microalgae.

**Figure. 8.**
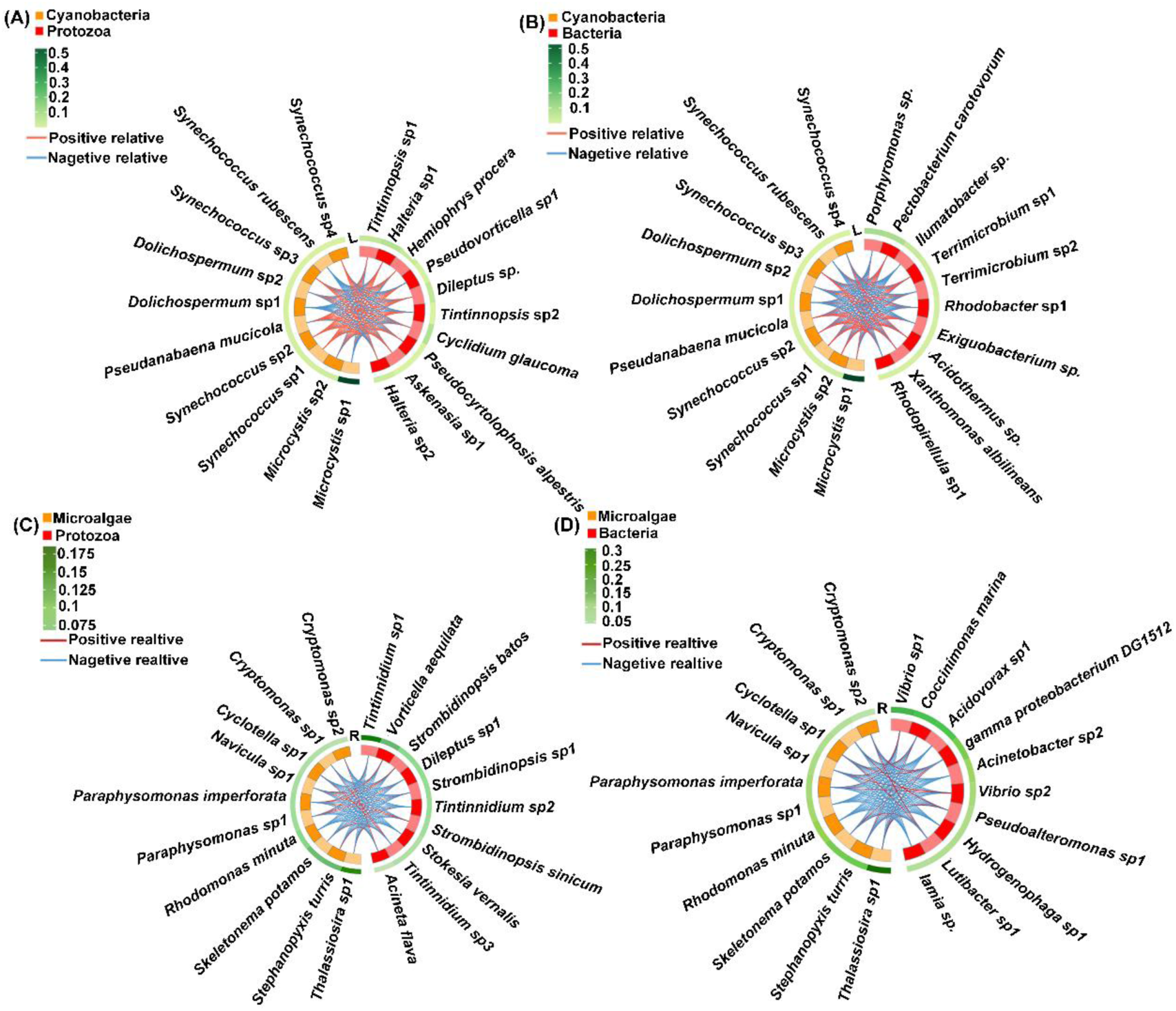
Correlation analysis of species level of microbial communities in lake and river (top10). Each column of the outer heat map represents a species, each layer represents a group, the grid color represents the average relative abundance of the species in the group, the middle line is the positive and negative correlation of the elements on both sides, the red line is the positive correlation, and the blue line is the negative correlation. **A and B.** On the left are cyanobacteria and on the right are the dominant species of other taxa in lake. **C and D.** On the left are eukaryotic microalgae and on the right are the dominant species of other taxa in river.

**Figure. 9.**
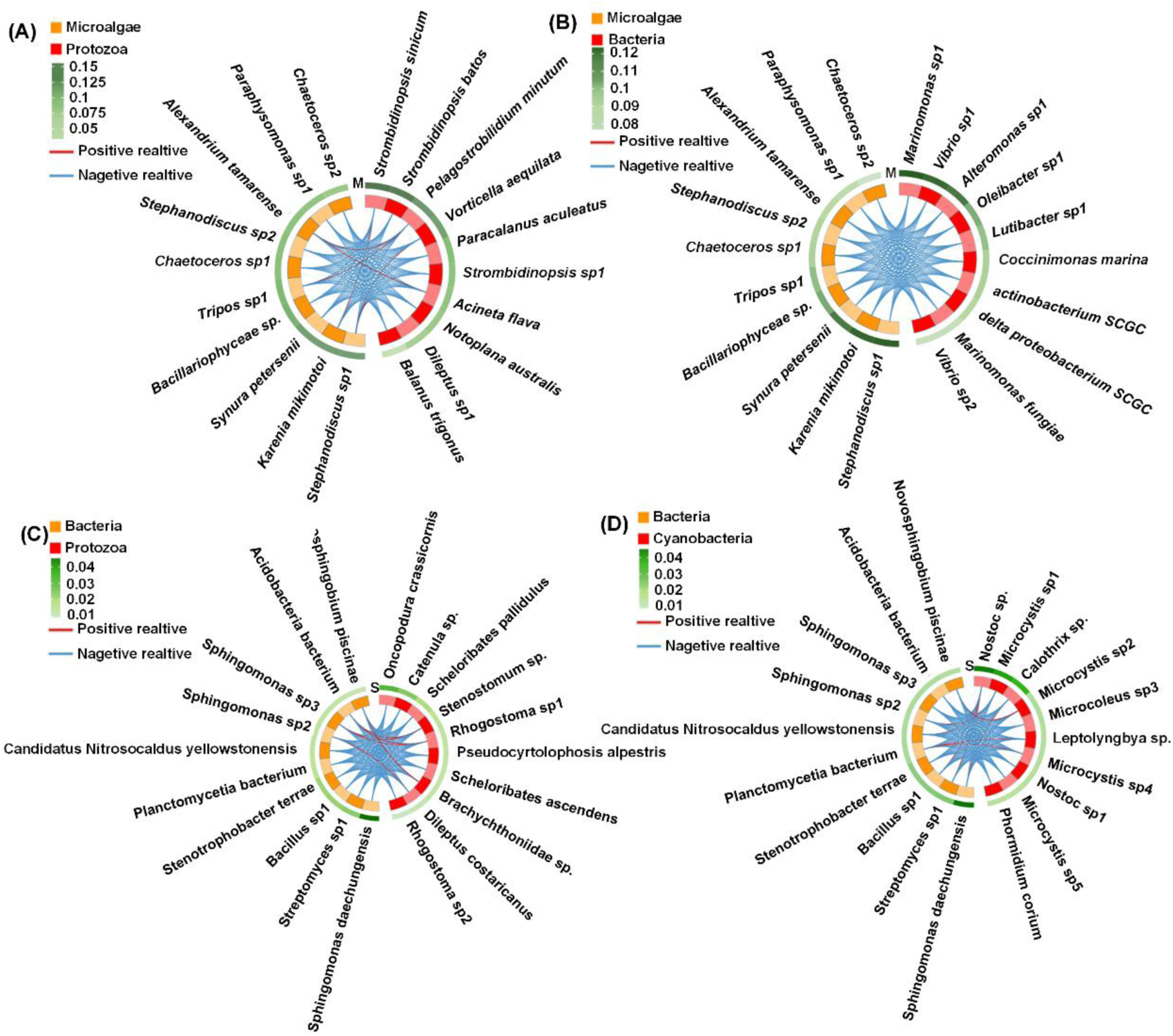
Correlation analysis of species level of microbial communities in marine and soil (top10). Each column of the outer heat map represents a species, each layer represents a group, the grid color represents the average relative abundance of the species in the group, the middle line is the positive and negative correlation of the elements on both sides, the red line is the positive correlation, and the blue line is the negative correlation. **A and B.** On the left are eukaryotic microalgae and on the right are the dominant species of other taxa in marine. **C and D.** On the left are bacteria and on the right are the dominant species of other taxa in soil.

Significant negative correlations were found among microalgae, cyanobacteria, protozoa and bacteria in the river (Fig. 8C, D and Fig. S1B). *Thalassiosira* sp1, the dominant species of microalgae with the highest mean relative abundance, was positively correlated with protozoa dominant species *Tintinnidium* sp3. (Fig. 8C). The three dominant *Tintinnidium* species of protozoa with higher relative abundance were mostly negatively correlated with other dominant microalgae species (Fig. 8C). The *Thalassiosira* sp1 was also positively correlated with the dominant species *gamma proteobacterium* DG1512 and *Lutibacter* sp1 of bacteria (Fig. 8E). The *Thalassiosira* sp1 was also positively correlated with the dominant species *Cylindrospermopsis* sp1 and *Dolichospermum affine* of cyanobacteria (Fig. S1B).

There were clear negative correlations among microalgae, cyanobacteria, protozoa and bacteria in marine environment (Fig. 9A, B and Fig. S1C). Only protozoa dominant species *Strombidinopsis sinicum* was positively correlated with microalgae dominant species *Karenia mikimotoi*. (Fig. 9A). While *Stephanodiscus* sp1 was negatively correlated with all dominant bacteria species and all dominant cyanobacteria species (Fig. 9B and Fig. S1C). There bacteria dominant species were significantly negatively correlated with all dominant species of protozoa, eukaryotic microalgae and cyanobacteria in soil (Fig. 9D, E and Fig. S1D). *Sphingomonas daechungensis*, the dominant bacterial species with the highest average relative abundance, was negatively correlated with protozoa and eukaryotic microalga dominant species (Fig. 9D and Fig. S1D), but positively correlated with the dominant cyanobacteria *Calothrix* sp. (Fig. 9E).

## Discussion

Currently, there are many research on diversity, dynamics and interaction of soil and water microorganisms (Aryal et al. 2015; Sigee 2005; Visser et al. 1992). However, a comprehensive comparative analysis of diversity, interactions and functional ecology of bacteria, cyanobacteria, eukaryotic microalgae and protozoa in soil and aquatic systems is lacking.

The community diversity difference of bacteria, cyanobacteria, microalgae and protozoa among soil, lake, river and marine environments was presented and compared in the schematic phylogenetic tree. Based on the taxonomy in Komárek et al (Komárek et al. 2014) ten orders were included in cyanobacteria from the ecosystem, in which Chroococcales, Nostocales, Oscillatoriales, Pleurocapsales and Synechococcales were revealed in our study from four environments. Alphaproteobacteria (Bacteria) was the most abundant bacteria in marine (Morris et al. 2002) while Actinobacteria was the most abundant bacteria in fresh water (Mehrshad et al. 2016). In our study, Alphaproteobacteria was the most abundant bacteria in lake, which may be related to interaction with cyanobacteria in lake since Proteobacteria can cause lysis of some cyanobacterial species (Eiler et al. 2004; Maruyama et al. 2003). Also, Alphaproteobacteria is the most abundant in soil (Spain et al. 2009).

Cercomonads (=Cercomonadida) (Protozoa) are tailed biflagellate heterotrophs found in soil, freshwater and marine ecosystems worldwide (Bass et al. 2009). In our study, Cercomonads were also found in soil and water, with relatively high abundance in water. Sessilida and Haptorida, belonging to ciliates, mainly inhabit in soil, fresh water, brackish water and marine environments (X Liu et al. 2012; Qu et al. 2022). Here we found that Sessilida had high mean relative abundance in aquatic environments, and Haptorida had high average relative abundance in freshwater environments and soil.

For microalgae, Chromulinales rarely found in previous reports were revealed in all of four environments here. Bacillariophyta are crucially important to the global ecosystem due to their role in regulating the world’s carbon and silicon cycles and their production of large amounts of organic material in aquatic environments (Bouchard 2021). Both Thalassiosirales and Naviculales are common diatoms. In previous studies, just several members of Thalassiosirales and Naviculales were studied from one environment (Maltsev et al. 2017; Martínez et al. 2023; Yu et al. 2022). Here we found that Thalassiosirales and Naviculales both exited in all environments of lake, soil, river and marine. Gymnodiniales, mostly existing in the marine (Reñé et al. 2015), were also just found in marine environment in this study.

From α diversity, it was showed that the species richness of cyanobacteria and bacteria in lake was both greater than that in other three environments. This may be related to the eutrophication and frequent occurrences of cyanobacterial blooms (including toxic species *Microcystis* sp.) throughout Taihu lake (Xu et al. 2017). The cyanobacteria and bacteria are closely interacted in water (Subashchandrabose et al. 2011). In our study, *Microcystis* sp. was also the dominant species of cyanobacteria in Taihu lake. The dynamics of *Microcystis* was significantly associated with the division of three major bacterial species, *Actinobacteria*, *Actinomyces* and *Proteobacteria* (L Liu et al. 2014) Thus, the bacteria diversity is also higher in lake. Similarly, the β diversity of cyanobacteria and bacteria also marked differences in lake from other environments. The species richness of protozoa and microalgae in aquatic environments were both greater than that in soil, which may be related with their close interaction in water. For example, the abundance and biomass of protozoa had significantly positive correlations with the biomass of Bacillariophyta and *Cryptophyta* in water (Sun et al. 2023). Similarly, there was a significant correlation between protozoan density and diatom density in freshwater environments (Kanavillil et al. 2013). This also can explain that part dominant microalgae in water were positively correlated with dominant protozoan species in this study (Fig. 7 and 8). However, protozoa and microalgae showed no significant differences in β diversity among different environments, suggesting that the species composition among different environments is generally similar except the dominant taxa.

For many years, the microbial food web interactions in water and soil both received a lot of attention (Gaedke et al. 2002). In aquatic environments, bacteria use organic compounds released into the environment by the death and decay of primary producers or other organisms to form a major granular food source for small and medium-sized heterotrophic protozoa (Fig.10) (OR Anderson 2001). This is consistent with the finding here that bacterial dominant species and protozoan dominant species were generally negatively correlated. In aquatic environment, microalgae and cyanobacteria provide carbon compounds and energy to bacteria and predators in the microfood web (Stockner et al. 1988). However, the algae blooms can also wipe out top-down predators and thus exacerbate the explosive spread of harmful algae (DM Anderson et al. 2021), which can explain the complex relationship between microalgae and protozoa dominant species in this study. In soil environment, the relationship between different microbes is also complicated. Among the diverse microbiota in the soil, microalgae and cyanobacteria are also the primary producers in the food chain that can grow in intimidating environments such as arid, semi-arid and wetland ecosystems (Abinandan et al. 2019). Terrestrial protozoa include heterotrophic flagellates, amoebae and ciliates (Adl 2003; Brussaard 1997), which prey on some bacteria and fungi. There are also predator-prey relationships between different protozoa (OR Anderson 2001). These are consistent with the complicated positive and negative relationships of various microbial dominant species in soil here. Anyway, our study provides the scientific basis for understanding the relationships of various microbial groups in the micro-food webs of different environments.

**Figure. 10.**
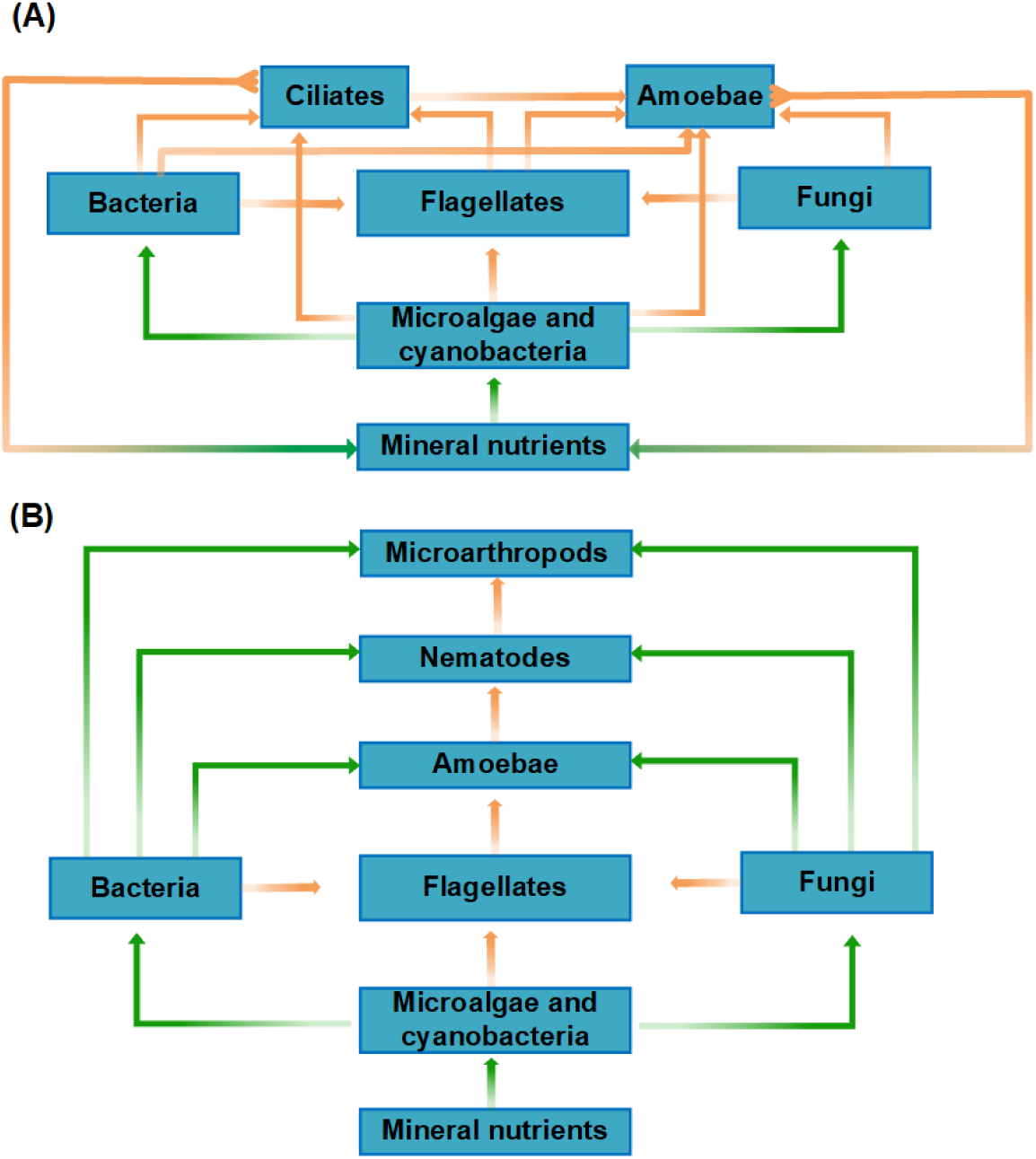
**A.** Food web based on different microbial communities in water environment. **B.** Food web based on different microbial communities in soil environment. The green arrow shows the flow of dissolved nutrients, and the orange arrow shows the relationship between predation and prey.

## Conclusion

Take Yangtze Delta of China as example, our study aimed to compare the community diversity and interaction of various microbes in micro-food chains of typical aquatic and soil environments. The results indicated that the diversity of bacteria, cyanobacteria, microalgae and protozoa among aquatic and soil environments is generally similar except the dominant taxa. The α diversity of cyanobacteria and bacteria in lake was higher than that in river, marine and soil. The β diversity of bacteria and cyanobacteria in lake was different from that in other environments, but all the environments showed no significant difference of β diversity for microalgae and protozoa. The cyanobacteria dominant species were mostly positively correlated with other microbial communities in lake. But the dominant species among bacteria, microalgae and protozoa were mostly negatively correlated in marine, river and soil.

## Acknowledgement

The bioinformatics analysis of the project was supported by the Bioinformatics Center of Nanjing Agricultural University.

## Ethical Approval

Not applicable

## Consent to Participate

The authors declare that the co-authors of the article all agreed to participate.

## Authors Contributions

Shanmei Zou designed the study and wirote the paper; Tiantian Sun conducted the experiment; Xinke Yu and Tiantian Sun colleted the samples; Lina Wei analyzed the data.

## Availability of data and materials

The NGS sequence data of all samples obtained in this study are deposited in NCBI. The SRA accession for the NGS sequences is PRJNA992270.

## References

Abinandan S, Subashchandrabose SR, Venkateswarlu K, Megharaj M (2019) Soil microalgae and cyanobacteria: the biotechnological potential in the maintenance of soil fertility and health. Critical reviews in biotechnology 39(8), 981–998.

Adl SM (2003) The ecology of soil decomposition: CABI.

Anderson DM, Fensin E, Gobler CJ, Hoeglund AE, Hubbard KA, Kulis DM, Landsberg JH, Lefebvre KA, Provoost P, Richlen ML (2021) Marine harmful algal blooms (HABs) in the United States: History, current status and future trends. Harmful Algae 102, 101975.

Anderson OR (2001) Protozoan ecology.

Archibald JM, Simpson AG, Slamovits CH, Margulis L, Melkonian M, Chapman DJ, Corliss JO (2017) Handbook of the Protists.

Aryal S, Karki G, Pandey S (2015) Microbial diversity in freshwater and marine environment. Nepal J. Biotechnol 3, 68–70.

Bass D, Howe AT, Mylnikov AP, Vickerman K, Chao EE, Smallbone JE, Snell J, Cabral Jr C, Cavalier-Smith T (2009) Phylogeny and classification of Cercomonadida (protozoa, Cercozoa): Cercomonas, Eocercomonas, Paracercomonas, and Cavernomonas gen. nov. Protist 160(4), 483–521.

Bolyen E, Rideout JR, Dillon MR, Bokulich NA, Abnet CC, Al-Ghalith GA, Alexander H, Alm EJ, Arumugam M, Asnicar F, Bai Y, Bisanz JE, Bittinger K, Brejnrod A, Brislawn CJ, Brown CT, Callahan BJ, Caraballo-Rodriguez AM, Chase J, Cope EK, Da Silva R, Diener C, Dorrestein PC, Douglas GM, Durall DM, Duvallet C, Edwardson CF, Ernst M, Estaki M, Fouquier J, Gauglitz JM, Gibbons SM, Gibson DL, Gonzalez A, Gorlick K, Guo J, Hillmann B, Holmes S, Holste H, Huttenhower C, Huttley GA, Janssen S, Jarmusch AK, Jiang LJ, Kaehler BD, Bin Kang K, Keefe CR, Keim P, Kelley ST, Knights D, Koester I, Kosciolek T, Kreps J, Langille MGI, Lee J, Ley R, Liu YX, Loftfield E, Lozupone C, Maher M, Marotz C, Martin BD, McDonald D, McIver LJ, Melnik AV, Metcalf JL, Morgan SC, Morton JT, Naimey AT, Navas-Molina JA, Nothias LF, Orchanian SB, Pearson T, Peoples SL, Petras D, Preuss ML, Pruesse E, Rasmussen LB, Rivers A, Robeson MS, Rosenthal P, Segata N, Shaffer M, Shiffer A, Sinha R, Song SJ, Spear JR, Swafford AD, Thompson LR, Torres PJ, Trinh P, Tripathi A, Turnbaugh PJ, Ul-Hasan S, van der Hooft JJJ, Vargas F, Vazquez-Baeza Y, Vogtmann E, von Hippel M, Walters W, Walters W, Wan Y, Wang M, Warren J, Weber KC, Williamson CHD, Willis AD, Xu ZZ, Zaneveld JR, Zhang YL, Zhu QY, Knight R, Caporaso JG (2019) Reproducible, interactive, scalable and extensible microbiome data science using QIIME 2 (vol 37, pg 852, 2019). Nat Biotechnol 37(9), 1091–1091. 10.1038/s41587-019-0252-6

Bouchard A. (2021). Incorporating Molecular Data in the Taxonomic Study of Diatoms: An Example Using Two Wellknown Genera, Frustulia and Navicula SS (Bacillariophyceae, Naviculales). Université d’Ottawa/University of Ottawa,

Brussaard L (1997) Biodiversity and ecosystem functioning in soil. Ambio, 563–570.

Das N, Chandran P (2011) Microbial degradation of petroleum hydrocarbon contaminants: an overview. Biotechnology research international 2011.

de Vargas C, Norris R, Zaninetti L, Gibb SW, Pawlowski J (1999) Molecular evidence of cryptic speciation in planktonic foraminifers and their relation to oceanic provinces. P Natl Acad Sci USA 96(6), 2864–2868. 10.1073/pnas.96.6.2864

Delgado-Baquerizo M, Oliverio AM, Brewer TE, Benavent-González A, Eldridge DJ, Bardgett RD, Maestre FT, Singh BK, Fierer N (2018) A global atlas of the dominant bacteria found in soil. Science 359(6373), 320–325.

Eiler A, Bertilsson S (2004) Composition of freshwater bacterial communities associated with cyanobacterial blooms in four Swedish lakes. Environmental Microbiology, 6(12), 1228–1243.

Endo H, Blanc-Mathieu R, Li Y, Salazar G, Henry N, Labadie K, de Vargas C, Sullivan MB, Bowler C, Wincker P (2020) Biogeography of marine giant viruses reveals their interplay with eukaryotes and ecological functions. Nature ecology & evolution 4(12), 1639–1649.

Fenchel T (2008) The microbial loop–25 years later. 366(1-2), 99–103.

Ficetola GF, Coissac E, Zundel S, Riaz T, Shehzad W, Bessière J, Taberlet P, Pompanon F (2010) An in silico approach for the evaluation of DNA barcodes. BMC Genomics 11(1), 1–10. 10.1111/1755-0998.12338

Fu KZ, Moe B, Li X, Le XC (2015) Cyanobacterial bloom dynamics in Lake Taihu. Journal of Environmental Sciences 32, 249–251.

Gaedke U, Hochstädter S, Straile D (2002) Interplay between energy limitation and nutritional deficiency: empirical data and food web models. Ecological Monographs 72(2), 251–270.

Gu C, Hu L, Zhang X, Wang X, Guo J (2011) Climate change and urbanization in the Yangtze River Delta. Habitat International 35(4), 544–552.

Guan B, An S, Gu B (2011) Assessment of ecosystem health during the past 40 years for Lake Taihu in the Yangtze River Delta, China. Limnology 12, 47–53.

Kanavillil N, Kurissery S (2013) Dynamics of grazing protozoa follow that of microalgae in natural biofilm communities. Hydrobiologia 718, 93–107.

Komárek J, Kaštovský J, Mareš J, Johansen JR (2014) Taxonomic classification of cyanoprokaryotes (cyanobacterial genera) 2014, using a polyphasic approach. Preslia 86(4), 295–335.

Klindworth, A., Pruesse, E., Schweer, T., Peplies, J., Quast, C., Horn, M., Glöckner, F.O., 2013. Evaluation of general 16S ribosomal RNA gene PCR primers for classical and next-generation sequencing-based diversity studies. Nucleic Acids Research 41, e1–e1. 10.1093/nar/gks808

Krzywinski M, Schein J, Birol I, Connors J, Gascoyne R, Horsman D, Jones SJ, Marra MA (2009) Circos: an information aesthetic for comparative genomics. Genome research 19(9), 1639–1645.

Leahy JG, Colwell RR (1990) Microbial degradation of hydrocarbons in the environment. Microbiological reviews 54(3), 305–315.

Liu L, Yang J, Lv H, Yu Z (2014) Synchronous dynamics and correlations between bacteria and phytoplankton in a subtropical drinking water reservoir. FEMS Microbiology Ecology 90(1), 126–138.

Liu X, Gong J (2012) Revealing the diversity and quantity of peritrich ciliates in environmental samples using specific primer-based PCR and quantitative PCR. Microbes Environments 27(4), 497–503.

Maltsev Y, Andreeva S, Kulikovskiy M, Podunaj J, Kociolek JP (2017) Molecular phylogeny of the diatom genus Envekadea (Bacillariophyceae, Naviculales). Nova Hedwigia 146, 241–252.

Martjnez LA, Sabbe K, Dasseville R, Daveloose I, Verstraete T, D’hondt S, Azémar F, Sossou AC, Tackx M, Maris T (2023) Long-term phytoplankton dynamics in the Zeeschelde estuary (Belgium) are driven by the interactive effects of de-eutrophication, altered hydrodynamics and extreme weather events. Sci Total Environ 860, 160402.

Maruyama T, Kato K, Yokoyama A, Tanaka T, Hiraishi A, Park H-D (2003) Dynamics of microcystin-degrading bacteria in mucilage of Microcystis. 46, 279–288.

Mehrshad M, Amoozegar MA, Ghai R, Shahzadeh Fazeli SA, Rodriguez-Valera F (2016) Genome reconstruction from metagenomic data sets reveals novel microbes in the brackish waters of the Caspian Sea. Appl Environ Micro 82(5), 1599–1612.

Mora C, Tittensor DP, Adl S, Simpson AG, Worm B (2011) How many species are there on Earth and in the ocean? PLoS biology 9(8), e1001127.

Morris RM, Rappé MS, Connon SA, Vergin KL, Siebold WA, Carlson CA, Giovannoni SJ (2002) SAR11 clade dominates ocean surface bacterioplankton communities. Nature 420(6917), 806–810.

Nübel, U., Garcia-Pichel, F., Muyzer, G., 1997. PCR primers to amplify 16S rRNA genes from cyanobacteria. Appl Environ Microbiol 63, 3327–3332. 10.1128/aem.63.8.3327-3332.1997.

Oksanen J, Blanchet F, Friendly M, Kindt R, Legendre P, McGlinn D, Minchin P, O’Hara R, Simpson G, Solymos P. (2019). vegan: Community Ecology Package. 2019. R package version 2.5-6. from Avilable online at https://CRAN.R-project.org/package=vegan (accessed …

Pan G, Zhang M-M, Chen H, Zou H, Yan H (2006) Removal of cyanobacterial blooms in Taihu Lake using local soils. I. Equilibrium and kinetic screening on the flocculation of Microcystis aeruginosa using commercially available clays and minerals. Environmental Pollution 141(2), 195–200.

Pinheiro J, Bates D, Debroy S, Sarkar D (2016) R Core Team.

Qu Z, Pan H, Gong J, Wang C, Filker S, Hu X (2022) Historical review of studies on cyrtophorian ciliates (Ciliophora, Cyrtophoria) from China. Microorganisms 10(7), 1325.

Reñé A, Camp J, Garcés E (2015) Diversity and phylogeny of Gymnodiniales (Dinophyceae) from the NW Mediterranean Sea revealed by a morphological and molecular approach. Protist 166(2), 234–263.

Sigee DC (2005) Freshwater microbiology: biodiversity and dynamic interactions of microorganisms in the aquatic environment: John Wiley & Sons.

Singer D, Seppey CV, Lentendu G, Dunthorn M, Bass D, Belbahri L, Blandenier Q, Debroas D, de Groot GA, De Vargas C (2021) Protist taxonomic and functional diversity in soil, freshwater and marine ecosystems. Environment International 146, 106262.

Spain AM, Krumholz LR, Elshahed MS (2009) Abundance, composition, diversity and novelty of soil Proteobacteria. The ISME journal 3(8), 992–1000.

Stahl D, Martinko J, Madigan M, Clark D. (2012). Brock biology of microorganisms. In: Benjamin-Cummings Pub Co., San Francisco.

Stockner JG, Porter KG. (1988). Microbial food webs in freshwater planktonic ecosystems. In Complex interactions in lake communities (pp. 69–83): Springer.

Subashchandrabose SR, Ramakrishnan B, Megharaj M, Venkateswarlu K, Naidu R (2011) Consortia of cyanobacteria/microalgae and bacteria: biotechnological potential. Biotechnology advances 29(6), 896–907.

Sun X, Zhang H, Wang Z, Huang T, Tian W, Huang H (2023) Responses of Zooplankton Community Pattern to Environmental Factors along the Salinity Gradient in a Seagoing River in Tianjin, China. Microorganisms 11(7), 1638.

Tedersoo L, Bahram M, Põlme S, Kõljalg U, Yorou NS, Wijesundera R, Ruiz LV, Vasco-Palacios AM, Thu PQ, Suija A (2014) Global diversity and geography of soil fungi. science 346(6213), 1256688.

Tortora GJ, Case CL, Bair III WB, Weber D, Funke BR (2004) Microbiology: an introduction.

Visser S, Parkinson D (1992) Soil biological criteria as indicators of soil quality: soil microorganisms. 7(1-2), 33–37.

Wang C, Wu S, Zhou S, Wang H, Li B, Chen H, Yu Y, Shi Y (2015) Polycyclic aromatic hydrocarbons in soils from urban to rural areas in Nanjing: Concentration, source, spatial distribution, and potential human health risk. Sci Total Environ 527, 375–383.

Wang H, Wu Q, Hu W, Huang B, Dong L, Liu G (2018) Using multi-medium factors analysis to assess heavy metal health risks along the Yangtze River in Nanjing, Southeast China. Environmental pollution 243, 1047–1056.

Wang J, Li S, Cui X, Li H, Qian X, Wang C, Sun Y (2016) Bioaccessibility, sources and health risk assessment of trace metals in urban park dust in Nanjing, Southeast China. Ecotoxicology and Environmental Safety 128, 161–170.

Wang M, Cao C, Li G, Singh RP (2015) Analysis of a severe prolonged regional haze episode in the Yangtze River Delta, China. Atmospheric Environment 102, 112–121.

Whitfield J (2005) Is everything everywhere? Science 310(5750), 960–961.

Xu H, Paerl HW, Zhu G, Qin B, Hall NS, Zhu M (2017) Long-term nutrient trends and harmful cyanobacterial bloom potential in hypertrophic Lake Taihu, China. Hydrobiologia 787, 229–242.

Ye W, Tan J, Liu X, Lin S, Pan J, Li D, Yang H (2011) Temporal variability of cyanobacterial populations in the water and sediment samples of Lake Taihu as determined by DGGE and real-time PCR. Harmful Algae 10(5), 472–479.

Yu P, Yang L, You Q, Bi Y, Wang Q (2022) A new freshwater species Conticribra sinica (Thalassiosirales, Bacillariophyta) from the lower reaches of the Yangtze River, China. Fottea 22(2), 238–255.

